# Drug-Induced p53 Activation Limits Pancreatic Cancer Initiation

**DOI:** 10.1101/2024.05.29.595146

**Authors:** Jennifer J Twardowski, Thomas I Heist, Zamira Guerra Soares, Emily S Berry, Luis I Ruffolo, Christoph Pröschel, Stephano S Mello

## Abstract

Pancreatic ductal adenocarcinoma (PDAC) is a highly lethal disease, initiated predominantly by mutations in *Kras*, which induce acinar-to-ductal metaplasia (ADM) and subsequent formation of precursor lesions, such as pancreatic intraepithelial neoplasia (PanIN). Progression to PDAC is frequently associated with mutations in the tumor suppressor *TP53*, presumably via disrupting p53-mediated cellular senescence of PanINs. Whether *TP53* also has tumor-suppressive activity in earlier phases of PDAC initiation has been less clear. In this study, we investigate the impact of pharmacological stabilization of the wild-type p53 protein on the formation of ADM in a *Kras*^*G12D*^-driven mouse model of PDAC. Our findings demonstrate that p53 stabilization via Nutlin-3a significantly reduces both ADM and PanIN formation by promoting the differentiation of ADM into acinar cells. This differentiation coincides with p53-dependent upregulation of the transcription factor Mist1 (*Bhlha15*), a critical inducer of acinar cell identity. Our results reveal a role for p53 in tissue repair and maintenance of homeostasis in tumor suppression and suggest pharmacological engagement of p53 as an intervention strategy to prevent PDAC initiation.

## Introduction

Pancreatic ductal adenocarcinoma (PDAC) is a deadly disease which is currently the third leading cause of cancer deaths^1^. Activating mutations in the proto-oncogene *Kras* are nearly ubiquitous in PDAC, detectable in over 90% of cases, followed by mutations of the tumor suppressor TP53, which affect 78% of PDAC cases^2^. The essential role of *Kras* mutations in the initiation and maintenance of PDAC has been established by several studies using genetically engineered mouse models^3-5^. On a cellular level, *Kras* mutations induce inflammation and dedifferentiation of normal pancreatic cells, leading to acinar-to-ductal metaplasia (ADM). During ADM, acinar cells lose their characteristic expression of Mist1 and Amylase and acquire ductal markers such as Sox9 and Cytokeratin-19^6-8^. This process ultimately results in the formation of preneoplastic pancreatic lesions known as pancreatic intraepithelial neoplasia (PanIN), which have the potential to develop into PDAC.

While *Kras* mutations are observed early and are considered driver mutations for PanIN formation, *TP53* mutations are typically found in more advanced PanIN, and are thought to promote the transition from PanIN to PDAC^9^. The p53 protein triggers various cellular responses by activating the transcription of target genes^10^. This transcriptional activation is a critical mechanism through which p53 exerts its tumor-suppressive effects^11, 12^. In the pancreas, p53-mediated tumor suppression has been shown to involve multiple p53 target genes, including *Cdkn1a* and *Ptpn14*. The *Cdkn1a* gene encodes p21, a p53-regulated cell cycle inhibitor that mediates oncogene-induced senescence in PanIN lesions^13, 14^. Similarly, *Ptpn14*, a protein tyrosine phosphatase, contributes to p53’s tumor-suppressive functions by decreasing proto-oncogenic YAP signaling in PDAC^12^. Whether p53 also plays a tumor-suppressive role during the early phases of pancreatic cancer initiation, particularly in the context of ADM formation, however, remains unclear.

A recent study using a hyperactive p53 mutant (*Trp53*^*F53Q,F54S*^, also referred to as *p53*^*53,54*^) indicated a strong reduction in ADM and PanIN formation in a mouse model for PDAC harboring a mutant *Kras* allele^15^, suggesting a possible role for p53 in limiting tumor initiation. Evidence that wild-type p53 can have a similar impact, however, is lacking. Here, we test the impact of drug-induced p53 stabilization on tumor initiation in an oncogenic *Kras*^*G12D*^-driven PDAC model. Our experiments indicate that pharmacological stabilization of wild-type p53 is sufficient to suppress initiation of *Kras*^*G12D*^-driven pancreatic cancer prior to PanIN formation, via a mechanism involving ADM-to-acinar cell differentiation and transactivation of a key driver of acinar cell identity.

## Results

### p53-dependent Nutlin-3a response reduces ADM and PanIN formation in the pancreas

To investigate the impact of p53 activation during the earliest stages of tumorigenesis in the pancreas, we used Nutlin-3a, a cis-imidazoline which inhibits Mdm2 (a negative regulator of p53^16^), in mice expressing mutant *Kras* in the pancreas (*Kras*^*LSL-G12D/+*^*;Ptf1a-Cre*, or *KC*)^17^. To accelerate PDAC initiation, we treated 2-month-old *KC* mice with cerulein to induce pancreatitis, followed by daily intraperitoneal injections of Nutlin-3a for one week, with DMSO as a vehicle control. Cerulein induces cell death, inflammation, and rapid ADM formation (**Figure 1a**), which resolves in a week through pancreatic regeneration in mice expressing wild-type Kras (**Figure S1a**)^18^. In contrast, in the presence of *Kras*^*G12D*^, pancreatic damage and ADM are not resolved^19^; instead, mutant Kras exacerbates ADM and drives the formation of PanIN lesions within one week (**Figure 1a-c**). Optimal Nutlin-3a dosing was determined by treating the animals with a range of drug concentrations (25, 12.5, 6, and 3 mg/kg of body weight) and assessing overall pancreatic histology and ADM formation using hematoxylin and eosin (H&E) staining and immunohistochemistry for the acinar cell marker Amylase. As expected, the pancreata of DMSO-treated *KC* mice lost most of their Amylase expression after a week, indicative of widespread ADM. In contrast, Nutlin-3a-treated *KC* pancreata, particularly those dosed with a low concentration of 6 mg/kg, presented with largely normal Amylase-expressing acinar cells (**Figure 1b**). To test whether Nutlin-3a may be preventing initial tissue injury, we analyzed pancreatic histology and Amylase expression in presence and absence of Nutlin-3a one and two days after the initial cerulein exposure. Loss of Amylase expression was observed to similar degrees in both conditions, indicating that Nutlin-3a does not mitigate the initial injury (**Figure S1b**). Our data thus demonstrate an unexpected effect of Nutlin-3a in promoting acinar cell regeneration in the presence of *Kras*^*G12D*^, with an optimal impact at a concentration of 6 mg/kg.

**Figure 1.**
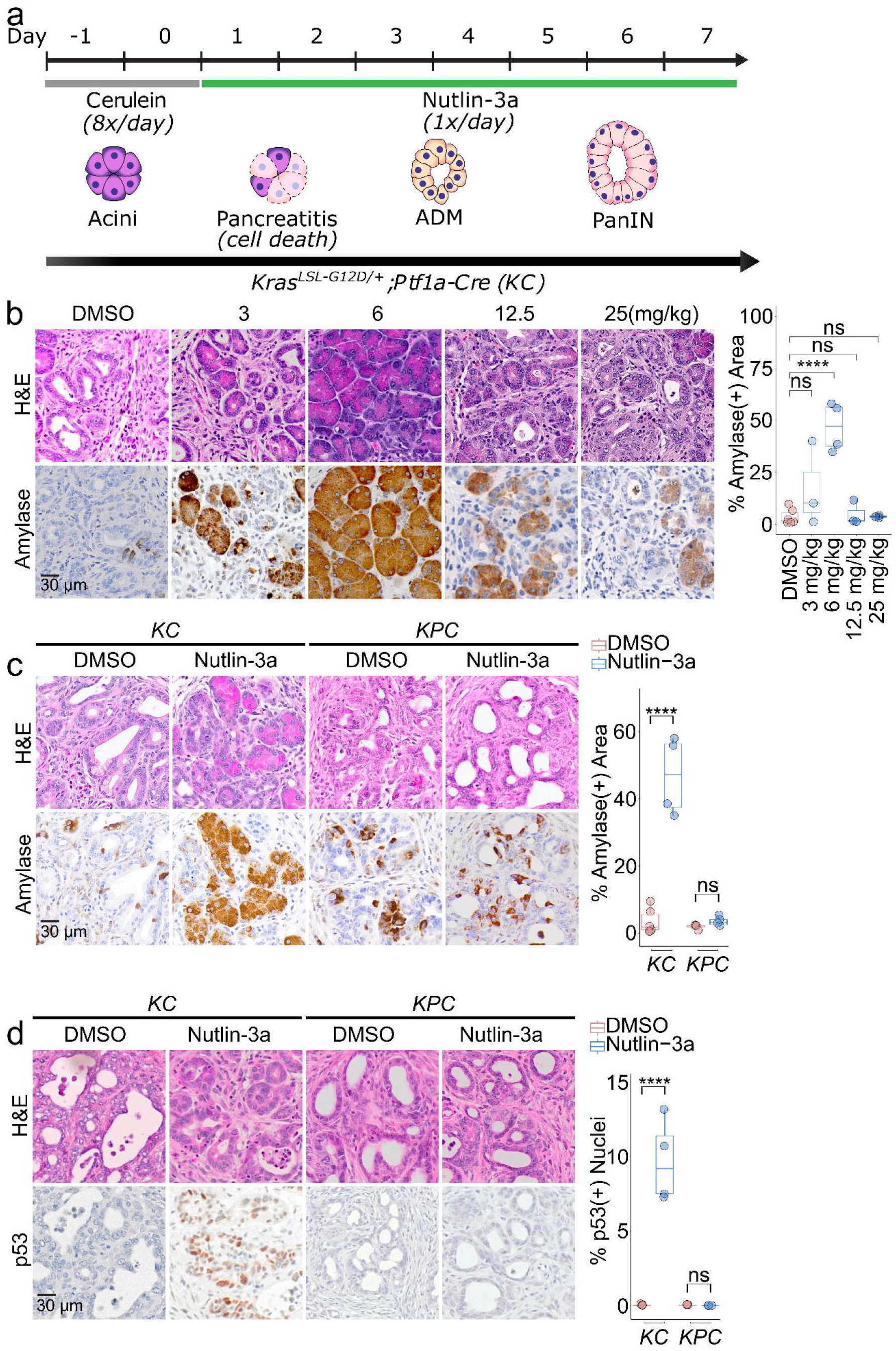
p53-dependent Nutlin-3a response reduces ADM formation in the pancreas. (A) Experimental timeline in which cerulein-induced pancreatitis is used to accelerate disease initiation, followed by treatment with either Nutlin-3a or DMSO (mock). (B) (Left) Representative images of H&E and immunostaining for Amylase in pancreatic tissue from *Kras*^*LSL-G12D/+*^*;Ptf1a-Cre* (*KC*) mice following cerulein treatment and administration of varying doses of Nutlin-3a or DMSO (mock) (counterstain: hematoxylin). (Right) Quantification of Amylase (+) area as a percentage of total pancreatic area (n=6 for mock-treated and n=3 or 4 for Nutlin-3a-treated *KC* mice). (C) (Left) Representative immunostaining for Amylase in the pancreata of *KC* and *Kras*^*LSL-G12D/+*^*;p53*^*fl/fl*^;*Ptf1a-Cre* (*KPC*) mice following one week of treatment with either 6 mg/kg of Nutlin-3a or DMSO (mock) (counterstain: hematoxylin). (Right) Quantification of Amylase (+) area as a percentage of total pancreas area (n=6 and 4 for mock-treated and Nutlin-3a-treated *KC* mice respectively, and n=4 for both mock-treated and Nutlin-3a-treated *KPC* cohorts). (D) (Left) Representative images of immunostaining for p53 in *KC* and *KPC* mice (counterstain: hematoxylin). (Right) Quantification of p53(+) nuclei as a percentage of total nuclei (n=6 and 4 for mock-treated and Nutlin-3a-treated *KC* mice respectively, and n=3 for both mock-treated and Nutlin-3a-treated *KPC* mice). Significance determined by unpaired two-tailed T test. (*): p<0.05; (**): p<0.01, (***): p<0.001.

To investigate whether the impact of Nutlin-3a on pancreatic regeneration is dependent on p53, we exposed 2-month-old mice expressing mutant Kras in the pancreas in absence of p53 (*Kras*^*LSL-G12D/+*^*;p53*^*fl/fl*^;*Ptf1a-Cre* or *KPC* mice)^20^ to cerulein and administered Nutlin-3a (6 mg/kg of body weight) as described above. In contrast to *KC* age-matched controls, *KPC* animals exhibited significant loss of acinar tissue that was not restored following treatment with Nutlin-3a, indicating a requirement of p53 for acinar regeneration (**Figure 1c**). We next tested whether a low dose of 6 mg/kg of body weight of Nutlin-3a is sufficient to activate p53. This was relevant, as previous *in vivo* experiments and clinical trials have utilized much higher concentrations^16, 21, 22^. As a surrogate for activation, we examined p53 expression in the pancreata of Nutlin-3a treated *KC* mice by immunohistochemistry and found increased p53 levels selectively in ADM structures, while no staining was seen in *KPC* control mice (**Figure 1d**). Selective p53 expression, alongside Mist1 and Amylase, is also found in pancreatic proto-acinar cells of wild-type mouse embryos at embryonic day 15.5 (**Figure S1c**), consistent with a regulation of p53 specific to cells committed to acinar identity. Furthermore, as expected for p53 activation, Nutlin-3a treatment reduced proliferation rates in both ADM and PanIN populations, as evidenced by reduced rates of Ki67 staining in *KC* but not in *KPC* mice exposed to Nutlin-3a (**Figure S1d-e**).

During ADM, acinar cells undergo metaplastic transformation and acquire ductal characteristics, eventually leading to the formation of PanINs^19^. This cellular reprogramming is driven by oncogenic mutations, such as *Kras*^*G12D*^, and involves the loss of acinar markers Mist1 and Amylase, as well as the gain of ductal markers such as Sox9 and Cytokeratin-19 ^23^. The formation of PanINs from ADM represents a critical early event in the initiation of PDAC. Given that p53 stabilization suppresses ADM, a precursor to PanIN formation, we predicted that Nutlin-3a would consequently impact PanIN formation. Following the same experimental approach described above, we investigated the impact of Nutlin-3a treatment on PanIN formation in the preneoplastic pancreas. PanIN lesions were identified using H&E, Alcian blue, and Mucin5ac, the latter of which are specific markers for PanIN^4, 24^. Alcian blue and Mucin5ac quantification revealed that Nutlin-3a-treated *KC* mice had a two-fold decrease in PanIN formation compared to mock-treated cohorts, suggesting that drug-induced p53 activation reduces PanIN formation. In contrast, PanINs were unaffected by Nutlin-3a treatment in *KPC* mice (**Figure 2**). In addition, Trichrome staining revealed a p53-dependent reduction in extracellular matrix remodeling and desmoplastic reaction in Nutlin-3a-treated *KC* mice (**Figure S2**), consistent with previous reports that established an association between desmoplastic reactions and PanIN lesions^25^. Thus, p53 stabilization via Nutlin-3a can suppress PanIN formation and inhibit cancer initiation in the pancreas.

**Figure 2.**
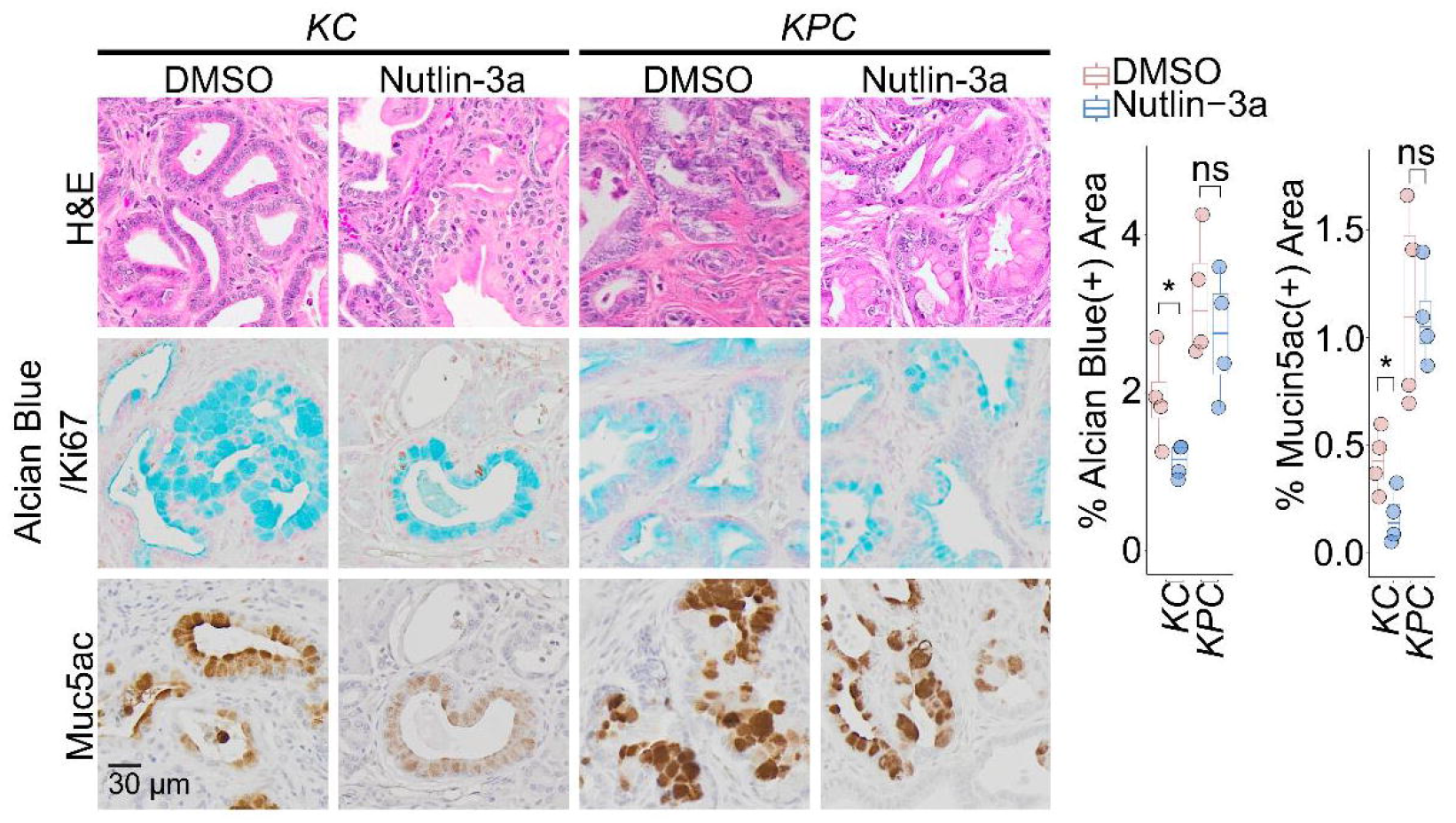
p53-dependent Nutlin-3a response reduces PanIN formation in the pancreas. (Left) Representative images of H&E and Alcian Blue staining (counterstain: Nuclear Fast Red), and immunostaining for Mucin5ac (counterstain: hematoxylin) in pancreatic tissue from cerulein-treated *KC* and *KPC* mice following one week of treatment with 6 mg/kg of Nutlin-3a or DMSO (mock). (Right) Quantification of Alcian Blue(+) or Mucin5ac(+) areas as a percentage of total pancreatic area (n=6 and 4 for mock-treated and Nutlin-3a-treated *KC* mice, and n=4 for both mock and Nutlin-3a-treated *KPC* mice).

### p53 promotes differentiation of ADM into acinar cells

Our findings suggest a function for p53 in promoting tissue repair through acinar cell regeneration. Given that Nutlin-3a treatment induced acinar regeneration in the presence of mutant Kras, we hypothesized that p53 activation supports the differentiation of ADM to acinar cells. To directly test this idea, we used a genetic lineage tracing mouse model with an inducible *Sox9-CreERT2* allele combined with *lox-stop-lox tdTomato*, a fluorescent reporter, allowing persistent labeling of ADM cells and their progeny. ADM cells exhibit ductal characteristics and express Sox9, thus enabling tdTomato activation following Cre-dependent recombination. *Kras*^*LSL-G12D/+*^*;Rosa26-CAG-LSL-tdTomato-WPRE;Sox9-Cre*^*ER*^ (*KTS*) mice were treated with tamoxifen and cerulein to induce mutant *Kras* expression as well as Cre activation resulting in ADM cell labeling. This allowed us to trace ADM cell fate after Nutlin-3a or DMSO (mock) treatment for one week (**Figure 3a**). Histological analysis of pancreata was performed to visualize tdTomato-positive ADM and acini. We found that in mock-treated animals, tdTomato-staining was detected largely in the ADM regions. In contrast, in Nutlin-3a treated animals, virtually all emerging acini had become tdTomato-positive (**Figure 3b**). Specificity and functionality of our Sox9-CreERT2-dependent model was tested in two distinct ways using *Rosa26-CAG-LSL-tdTomato-WPRE;Sox9-Cre*^*ER*^ (*TS*) mice. First, we confirmed that expression of Cre recombinase was highly specific, as no tdTomato-positive cells were observed in the absence of tamoxifen (**Figure S3a**). Second, we validated that lox-stop-lox tdTomato recombination occurred in in Sox9-expressing normal pancreatic ducts, but not in acinar cells lacking Sox9 expression, confirming that recombination took place selectively in Sox9-expressing cells (**Figure S3b)**. Taken together, these results strongly support the conclusion that Nutlin-3a-mediated p53 activation promotes the differentiation of ADM into acini in the context of *Kras*^*G12D*^, presumably preventing progression to PanIN (**Figure 3c**).

**Figure 3.**
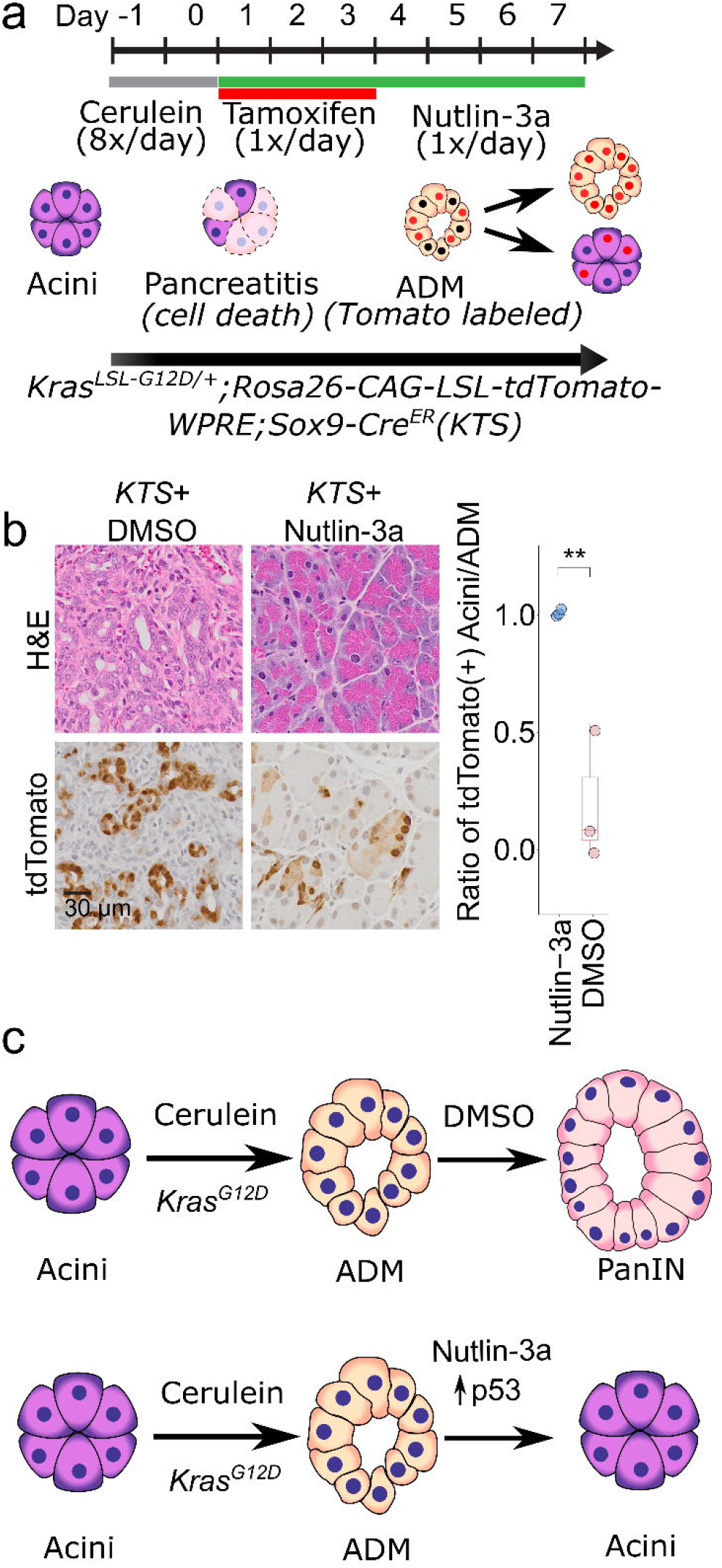
p53 promotes differentiation of ADM into acinar cells. (A) Experimental timeline depicting cerulein-induced pancreatitis followed by tamoxifen administration and treatment with either Nutlin-3a or DMSO (mock). (B) (Left) Representative H&E images and immunostaining for tdTomato in ADM and acini from Nutlin-3a and DMSO (mock) treated *Kras*^*LSL-G12D/+*^*;Rosa26-CAG-LSL-tdTomato-WPRE;Sox9-Cre*^*ER*^ (*KTS*) mice. (Right) Quantification of tdTomato(+) acini as a ratio of total tdTomato(+) ADM and acini in *KTS* mice (n=3 for both mock and Nutlin-3a-treated *KTS* mice). (C) Proposed model for Nutlin-3a impact on resolving pancreatic damage during cerulein-induced pancreatitis in the context of mutant Kras. Significance determined by unpaired two-tailed T test. (**): p<0.01.

### p53 controls the expression of acinar cell identity genes

Having established that p53 activation can promote ADM-to-acinar cell differentiation and suppress PanIN formation in the presence of mutant Kras, we sought to elucidate underlying mechanisms by which p53 can impact acinar cell fate. For this purpose we performed gene set enrichment analysis (GSEA) using RNA-seq data derived from sorted pancreatic p53-proficient and p53-deficient precursor lesions^12^ and cross-compared the results with three ChIP-seq data sets specific for key acinar cell identity transcription factors, Mist1, Ptf1a and Rbpjl^26, 27^, respectively. This revealed a significant overlap between genes regulated by p53 and Mist1, while such a relationship was not detected for Ptf1a or Rbpjl (**Figure 4a, Figure S4a**). Further, interrogation of p53 ChIP-seq data derived from murine and human fibroblasts^28, 29^ revealed p53-binding to both murine and human genes encoding Mist1, i.e. *Bhlha15* at multiple sites (**Figure 4b**), as seen for other p53 target genes in various cell types^30^. Notably, *Bhlha15* expression was significantly decreased in the absence of p53 in preneoplastic pancreatic cells, consistent with the idea that *Bhlha15* levels in the pancreas are sensitive to p53 (**Figure 4c**). Conversely, we observed Mist1 expression in ADM regions of Nutlin-3a-treated *KC* mice, while no Mist1 staining was present in the ADM of mock-treated mice, suggesting that Mist1 expression, and thus acinar cell identity was being driven by p53 activation (**Figure 4d**). In addition, *Bhlha15* expression is also sensitive to p53 in murine and human fibroblasts treated with the DNA damaging agent doxorubicin, (**Figure 4e-f; Figure S4b**), suggesting that p53 can transactivate *Bhlha15*, presumably directly, across distinct cell types and in different contexts, such as oncogenic stress and DNA damage.

**Figure 4.**
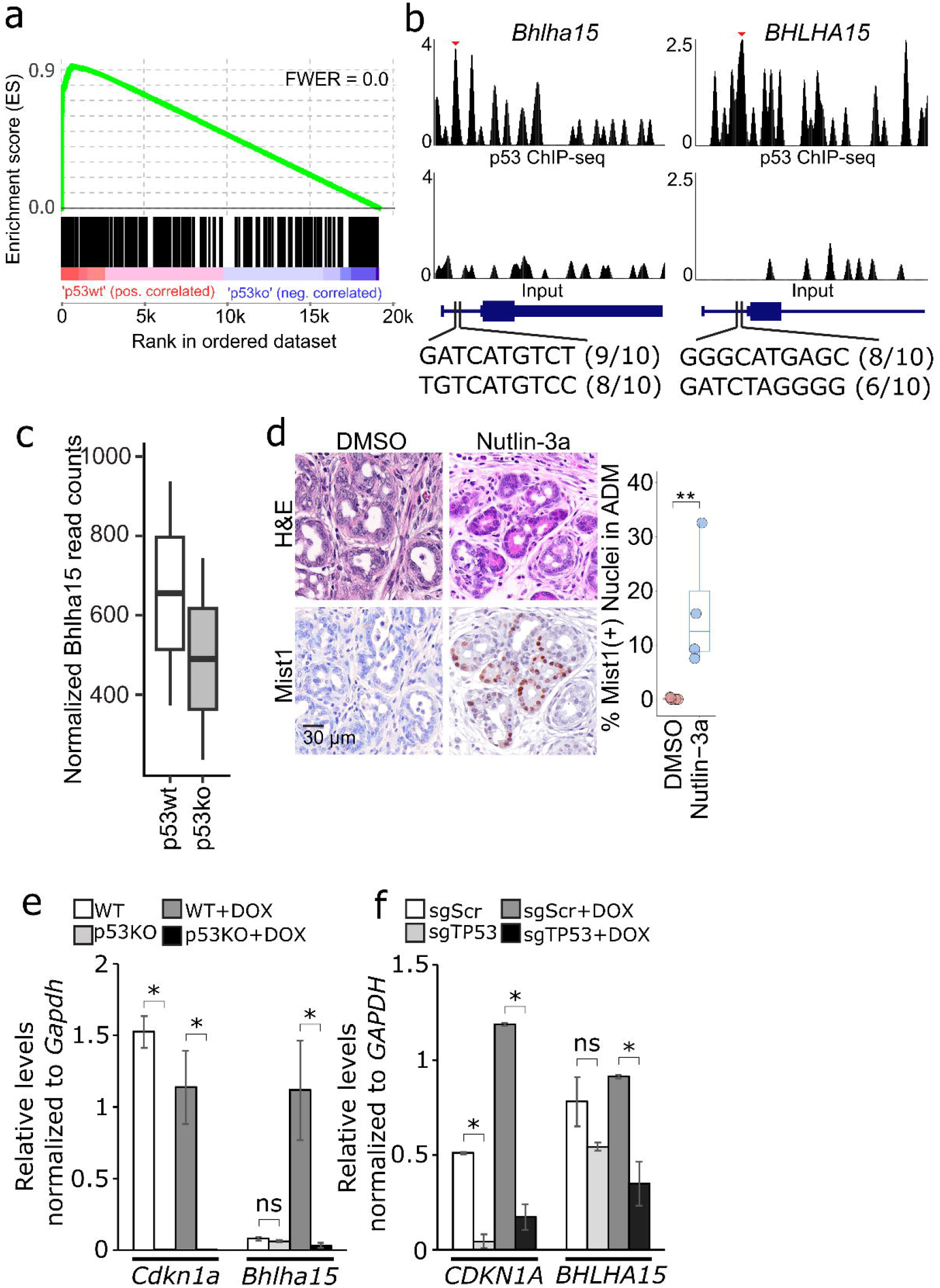
p53 controls the expression of acinar cell identity genes. (A) GSEA showing RNA-seq data from sorted p53 wild-type and p53 knockout pancreatic precursor lesions cross-compared with Mist1 ChIP-seq data. p53 ChIP-Seq peaks at the *Bhlha15/ BHLHA15* gene in mouse and human fibroblasts. Numbers in parentheses indicate the numbers of base pairs in individual half-sites matching the consensus p53 response element sequence. Number between brackets indicate the space in nucleotides (nts) between the two half-sites. Expression of *Bhlha15* in RNA-seq data from sorted p53 wild-type and p53 knockout pancreatic precursor lesions. (D) (Left) Representative H&E images and immunostaining for Mist1 in pancreatic sections from *KC* mice (counterstain: hematoxylin). (Right) Quantification of Mist1(+) nuclei within ADM (n=6 and 4 for mock and Nutlin-3a-treated mice respectively). (E) RT-qPCR analysis of *Cdkn1a* and *Bhlha15* in wild-type and p53-null MEFs treated with 0.2µg/mL of doxorubicin (DOX) for 8 hours; normalized to Gapdh. (F) RT-qPCR analysis of *CDKN1A* and *BHLHA15* in human fibroblasts following CRISPRi-mediated p53 knockdown (sgScr: non-targeting scrambled guide; and sgTP53: guide targeting TP53 gene) and treatment with DOX for 8 hours; normalized to GAPDH. Significance determined by unpaired two-tailed T test. (*): p<0.05; (**): p<0.01.

## Discussion

Our study reveals that activation of wild type p53 can reverse early tissue damage associated with tumor initiation in a *Kras*^*G12D*^-driven mouse model for pancreatic ductal adenocarcinoma (PDAC). We provide compelling evidence that pharmacological stabilization of endogenous wild-type p53 protein can effectively limit ADM and PanIN formation in the pancreas following p53-mediated ADM-to-acinar differentiation. This involves p53-induced upregulation of *Bhlha15*, a gene encoding the transcription factor Mist1, which is essential for acinar cell differentiation. Our findings suggest that engaging p53 activity during oncogenic stress can lead to tumor suppression by means of acinar cell differentiation and tissue repair.

Mechanisms of p53-mediated tumor suppression are highly diverse. In addition to cell cycle arrest and apoptosis, they also involve induction of senescence and reprogramming of cell metabolism controlled by a variety of p53 target genes^11, 31, 32^. In this study, we show that engagement of wild-type p53 via pharmacological activation can repair pancreatic damage caused by cerulein and *Kras*^*G12D*^ that otherwise appears irreversible. Notably, in this context, p53 activation stimulates the differentiation of ADM into acini, presumably via induction of Mist1. This is reminiscent of observations in lung indicating that p53 activation can promote differentiation of AT1 alveolar cells through direct upregulation of AT1-related genes *Pdgfa, Fbln5* and *Fam174b*^33, 34^. Previous work suggests that p53’s major role in PDAC suppression relates to its ability to induce senescence in PanIN lesions^13, 14^. In contrast, our work has focused on ADM as the earliest indicator of tissue damage mediated by *Kras*^*G12D*^ in the pancreas, thus revealing an additional role of p53 in tumor suppression involving tissue repair. In this context, it is worth noting that p53 knockout mice with wild-type *Kras* do not display overt symptoms of acinar hypoplasia. Thus, while not essential for acinar cell differentiation, p53 appears to play a critical role in supporting pancreatic tissue differentiation and integrity in the context of oncogenic stress.

The p53 tumor suppressor may engage several cellular processes simultaneously to elicit its full impact^35^. Given the crucial role of Mist1 in maintaining acinar identity^26, 36^, its upregulation in ADM cells following Nutlin-3a treatment suggests that Mist1 may play a significant role in p53-mediated pancreatic regeneration. In addition, the p53-regulated gene *Neat1*, encoding a long non-coding RNA, has been found to promote the expression of pro-differentiation programs, thereby reducing the risk of malignant transformation and preserving normal pancreas cell identity in the context of mutant *Kras*^37^. Our data presented here also indicate an inhibition of ADM cell proliferation rates in response to Nutlin-3a exposure, suggesting that a reduced rate of cell cycle progression may help facilitate acinar differentiation, as previously reported^38^. Collectively, these observations highlight the complexity of p53-regulated cellular processes in pancreatic regeneration.

In summary, our study demonstrates that pharmacological activation of wild-type p53 can promote pancreatic tissue repair in response to oncogenic challenge. In this process, differentiation of ADM into acini plays a significant role in mitigating the emergence of preneoplastic lesions. Harnessing p53-mediated pathways for therapeutic strategies may thus involve pharmacological activation of p53 to promote metaplastic cell differentiation as a preventative PDAC intervention.

## Materials and Methods

### Animal Models

All animal experiments were previously approved and performed according to the University Committee on Animal Resources at the University of Rochester (UCAR). Mouse models for PDAC harboring a mutant *Kras* allele in the pancreas were generated by crossing *Kras*^*LSL-G12D/+*^ mice^39^ with *Ptf1a-Cre* mice^40^. Mouse models for PDAC without p53 were generated by crossing *p53*^*fl/fl*^ mice^20^ to *Kras*^*LSL-G12D/+*^*;Ptf1a-Cre* (*KC*) mice. To label ADM cells and investigate the fate of metaplastic acini, *Rosa26-CAG-LSL-tdTomato-WPRE* mice^41^ were bred to *Sox9-Cre*^*ER*^ mice^42^, allowing the induction of Tomato specifically in Sox9-positive cells. *LSL-tdTomato-WPRE;Sox9-Cre*^*ER*^ (*TS*) mice were further bred to *Kras*^*LSL-G12D/+*^ mice. Induction of Sox9-Cre was performed by the treatment with tamoxifen (MP Biomedicals 156738) dissolved in corn oil and ethanol and administered via oral gavage once daily for three days as previously described^43^.

To induce acute pancreatitis, 2-month-old mice were administered eight hourly intraperitoneal injections of 10 µg/mL cerulein (Bachem 4030451) in PBS over the course of two days, at a dosage of 100 µg/kg of body weight, as previously described^44^. For drug-induced p53 activation experiments, mice were given intraperitoneal injections of either Nutlin-3a (APExBIO A3671) diluted in DMSO or DMSO alone (mock treatment) once daily for one week, at dosages of 3 mg/kg, 6 mg/kg, 12.5 mg/kg, and 25 mg/kg.

### Immunohistochemistry

After experimental treatments, mouse pancreata were extracted and stored in 10% formalin solution. Tissue samples were processed, paraffinized, and sectioned by the Wilmot Cancer Institute Histopathology Core and Histology, Biochemistry, and Molecular Imaging (HBMI) Core. Immunohistochemistry was performed on formalin-fixed, paraffin-embedded tissue sections according to a standard protocol. Primary antibodies included mouse monoclonal anti-AMYLASE (Santa Cruz Biotechnology sc-46657, 1:100), rabbit monoclonal anti-MIST1 (Cell Signaling Technology 14896T, 1:100), rabbit polyclonal anti-p53 (Invitrogen PA5-88098, 1:300), mouse monoclonal anti-KI67 (BD Pharmingen 550609, 1:100), and rabbit polyclonal anti-RFP (Rockland 600-401-379, 1:100). Secondary antibodies included biotinylated goat anti-rabbit (Vector Laboratories BA-1000, 1:200) and anti-mouse (Santa Cruz Biotechnologies sc-516142, 1:200) IgG antibodies, Alexa Fluor™ 555 goat anti-mouse (Invitrogen A28180, 1:200), and Alexa Fluor™ 647 donkey anti-rabbit (Invitrogen A-31573, 1:200) antibodies. Binding avidity for biotinylated antibodies was increased using the Vectastain ABC Kit (Vector Laboratories PK-4000). Binding of biotinylated antibodies was visualized using a DAB peroxidase substrate kit (Vector Laboratories SK-4100). Counterstains included Hematoxylin QS (Vector Laboratories H-3404), DAPI (eBioscience 00-4959-52), and Nuclear Fast Red solution (IHC World IW-3000B). Other stains included Alcian blue solution (IHC World IW-3000A), eosin, and Masson’s trichrome (provided by HBMI Core).

### Tissue Analysis

Stained tissue samples were scanned using an Olympus VS120 Virtual Slide Microscope and Scanner. Image analysis and quantification of IHC markers were performed using Visiopharm (v.2019.07; Hørsholm, Denmark) and QuPath^45^. For cytoplasmic staining, positively stained area was quantified and presented as a percentage of total pancreatic area. For nuclear staining, positively stained nuclei were counted and presented as a percentage of total nuclei. For nuclear quantification within metaplastic cells, ADM were manually set as regions of interest (ROI) based on morphology, and nuclei within the ROI were quantified. For nuclear quantification within PanIN lesions, lesions were set as ROI using Alcian blue positivity, and nuclei within the ROI were quantified. A Shapiro-Wilk test was used to evaluate distribution of the data. For normally distributed samples, an unpaired 2-tailed Student’s T-test was performed. For data not normally distributed, a Kruskal-Wallis test was performed. Significant outliers were excluded. Statistical tests were performed using RStudio. P-values less than 0.05 were considered statistically significant.

### Sequencing Data and Analysis

p53 ChIP-seq data in mouse and human cells used for the identification of novel p53 target genes were previously published^28, 29^ and obtained from Gene Expression Omnibus (GEO, GSE46240 and GSE55727 respectively). RNA-seq data from preneoplastic lesions sourced from p53 wild-type (*Kras*^*LSL-G12D/+*^*;Pdx1-Cre*) and p53 knockout (*Kras*^*LSL-G12D/+*^*;Pdx1-Cre;p53*^*-/-*^) mice were also previously published^12^ and obtained from GEO (GSE94566). ChIP-seq data was aligned to the mouse genome (mm10) or human genome (hg38) using Bowtie2^46^. Peaks were called using MACS2^47^ and bigwig files for visualization of p53 ChIP-seq peaks were generated using deepTools^48^. ChIPseeker^49^ was used to annotate p53 ChIP-seq peaks to genes. RNA-seq data was aligned using Salmon^50^ and analyzed using DESEQ2^51^. Genes with a false discovery rate (FDR) of 0.1 or less were considered significant. Gene set enrichment analysis (GSEA)^52^ was conducted using the standard mode and the RNA-seq data from GSE94566, which includes pancreatic preneoplastic lesions from both p53 wild-type and p53 knockout mice. The analysis utilized the top 250 genes associated with the most significant peaks from previously published ChIP-seq data for Mist1, Ptf1a, and Rbpjl in pancreatic cells (GSE86289 and GSE47459) ^26, 27^ as signatures.

### Cell Culture

Mouse embryonic fibroblasts (MEFs) were obtained from *p53*^*fl/fl*^ mice and were infected with Ad5 CMV-Cre or Ad5 CMV-empty adenovirus to obtain p53 knockout and p53 wild-type cells. Telomerase-immortalized MRC5 cells were a kind gift from Dr. Joshua Munger^53^. Both cells were maintained in DMEM with 10% FBS. CRISPR interference (CRISPRi) was performed in MRC5 cells using a previously validated^54^ guide RNA against p53 (GCCCTACGCCCAGCACCGGG) or a non-targeting scrambled guide (GCTGCATGGGGCGCGAATCA). Doxorubicin (Sigma) was used at 0.2 μg/mL to cause DNA damage and activate p53 in both mouse and human cells.

### RT-qPCR analysis

For RT–qPCR, RNA was isolated using Trizol reagent (Invitrogen) and reverse-transcribed using High-Capacity cDNA Reverse Transcription Kit (Applied Biosystems) and random primers. PCR was performed in triplicate using PerfeCTa SYBR Green FastMix (Quantabio) and a QuantStudio 5 Real-Time PCR machine (Applied Biosystems), and the results were computed relative to a standard curve made with cDNA pooled from all samples. The average of three independent experiments were plotted and the primer sequences for RT–qPCR are listed in the **Supplemental Table 1**.

## Acknowledgements

The authors would like to thank Dr. Joshua Munger for providing immortalized MRC5 cells. We would also like to thank Jeffrey Fox and Vidya Venkatramani in the CMSR Histology Core for providing tissue processing and sectioning services. Lastly, we would like to thank Dr. Hartmut “Hucky” Land for his invaluable feedback throughout the creation of this work.

## Appendix

**Figure S1:** (A) Representative images of H&E-stained pancreata from *Kras*^*WT*^ mice one and seven days following induction of acute pancreatitis. (B) Representative images of IHC staining for Amylase in Nutlin-3a-treated and mock-treated *KC* mice one and two days after induction of acute pancreatitis (counterstain: hematoxylin). (C) Representative images of IHC staining for p53, Mist1, and Amylase in embryonic mouse pancreatic tissue (counterstain: hematoxylin). (D) (Left) Representative images of IHC staining for Ki67 in *KC* and *KPC* mice (counterstains: Alcian blue and Nuclear Fast Red). (Right) Quantification of Ki67(+) nuclei within selected PanIN. n=6 mock-treated *KC*; n=4 Nutlin-3a-treated *KC*, mock-treated *KPC*, and Nutlin-3a-treated *KPC* mice. (E) (Left) Representative images of IHC staining for Ki67 in *KC* and *KPC* mice (counterstain: hematoxylin). (Right) Quantification of Ki67(+) nuclei within ADM. n=6 mock-treated *KC*; n=4 Nutlin-3a-treated *KC*, mock-treated *KPC*, and Nutlin-3a-treated *KPC* mice. Significance determined by unpaired two-tailed T test. (*): p<0.05; (**): p<0.01.

**Figure S2:** (Left) Representative images of Trichrome staining in *KC* and *KPC* mice. (Right) Quantification of fibrotic tissue as a percentage of total pancreatic area. n=6 mock-treated *KC*; n=4 Nutlin-3a-treated *KC*, mock-treated *KPC*, and Nutlin-3a-treated *KPC* mice. Significance determined by unpaired two-tailed T test. (*): p<0.05.

**Figure S3:** (A) Representative images of *Rosa26-CAG-LSL-tdTomato-WPRE;Sox9-Cre*^*ER*^ (*TS*) and *Kras*^*LSL-G12D/+*^*;Rosa26-CAG-LSL-tdTomato-WPRE;Sox9-Cre*^*ER*^ (*KTS)* mouse pancreata after cerulein-induced pancreatitis, but no treatment of tamoxifen. Examples of ducts and acini are shown. (B) Examples of tdTomato(+) ducts in *TS* mice, as indicated by H&E staining, IHC for tdTomato (counterstain: hematoxylin), and IF for RFP and Amylase (counterstain: DAPI). (C) Examples of tdTomato(+) acini in *TS* mice, as indicated by H&E staining, IHC for tdTomato (counterstain: hematoxylin), and IF for RFP and Amylase (counterstain: DAPI).

**Figure S4:** (A) GSEA using RNA-seq data from sorted *p53*^*WT*^ and *p53*^*KO*^ pancreatic precursor lesions compared to Ptf1a (left) and Rbpjl (right) ChIP-seq data. (B) RT-qPCR verification of CRISPRi knockdown of p53.

**Table S1:** Sequences of primers used in qPCRs.

