## Supplemental Figures for "Drug-Induced p53 Activation Limits Pancreatic Cancer Initiation"

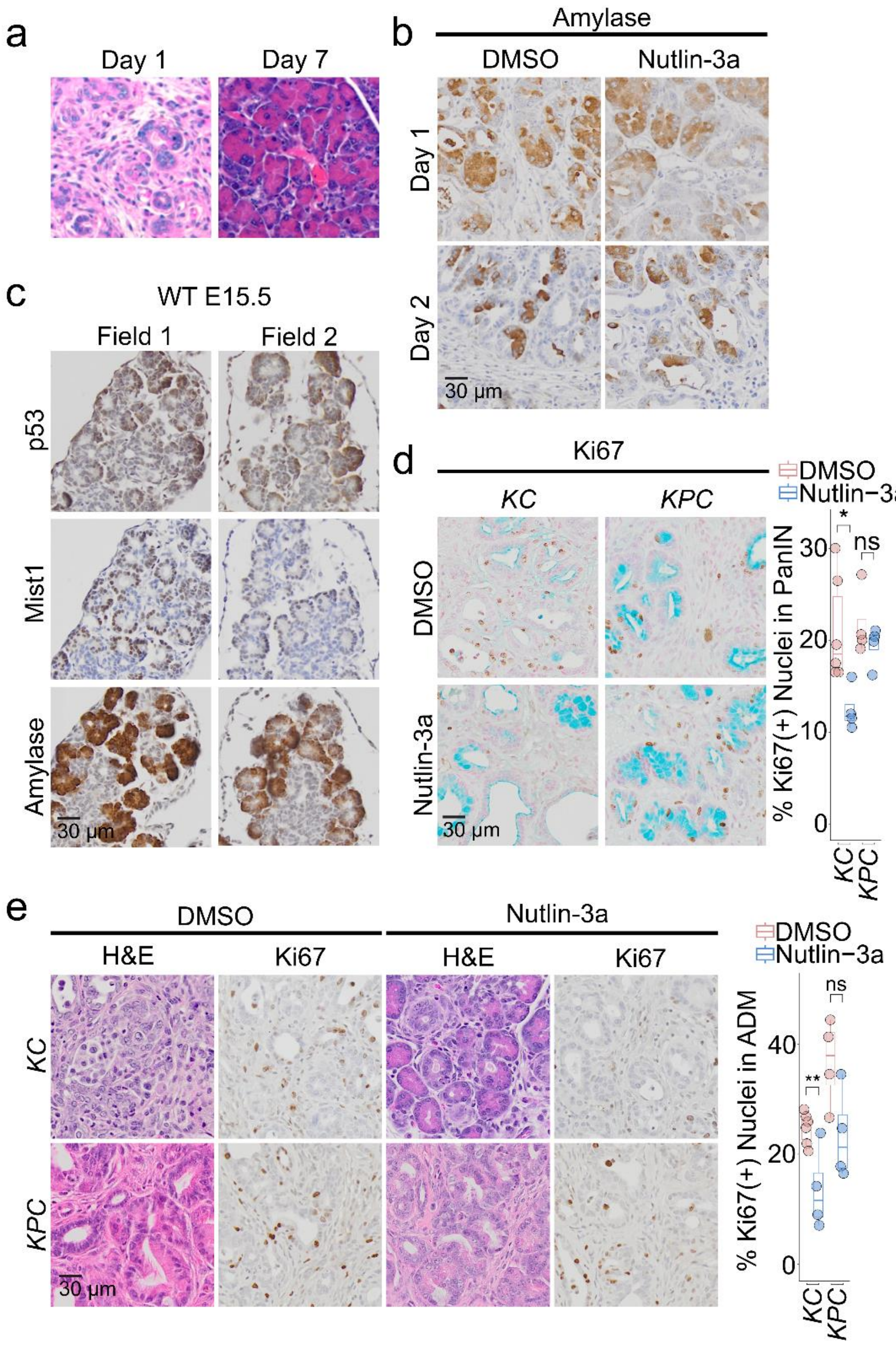


**Figure S1:**

a. Representative H&E images of wild-type Kras mice treated with cerulein, at day 1 (left) and day 7 (right) after treatment.

b. Representative images of IHC staining for Amylase in Nutlin-3a-treated and mock-treated *KC* mice one and two days after induction of acute pancreatitis (counterstain: hematoxylin).

c. Representative images of IHC staining for p53, Mist1, and Amylase in embryonic mouse pancreatic tissue (counterstain: hematoxylin).

d. (Left) Representative images of IHC staining for Ki67 in *KC* and *KPC* mice (counterstains: Alcian blue and Nuclear Fast Red). (Right) Quantification of Ki67(+) nuclei within selected PanIN. n=6 mock-treated *KC*; n=4 Nutlin-3a-treated *KC*, mock-treated *KPC*, and Nutlin-3a-treated *KPC* mice.

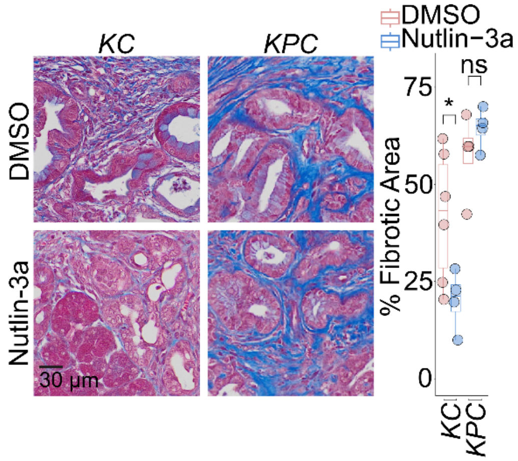


**Figure S2:**

(Left) Representative images of Trichrome staining in *KC* and *KPC* mice. (Right) Quantification of fibrotic tissue as a percentage of total pancreatic area. n=6 mock-treated KC; n=4 Nutlin-3a-treated *KC*, mocktreated *KPC*, and Nutlin-3a-treated *KPC* mice. Significance determined by unpaired two-tailed T test. (*): $p<0.05$.

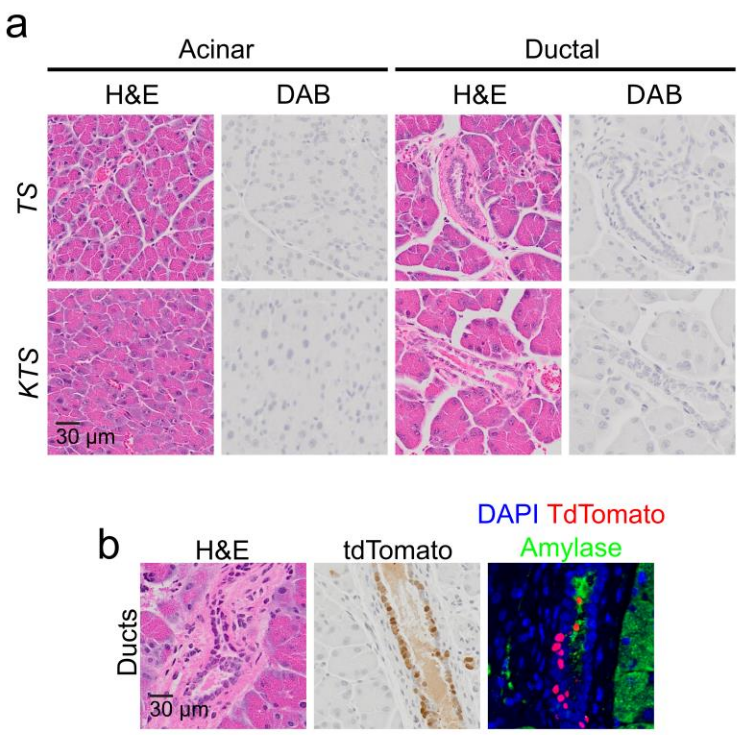


**Figure S3:**

a. Representative images of pancreata from *Rosa26-CAG-LSL-tdTomatoWPRE;Sox9-CreER (TS)* and *Kras*$^{LSL\text{-}G12D/+}$*;Rosa26-CAG-LSL-tdTomatoWPRE;Sox9-CreER (KTS)* mice after cerulein-induced pancreatitis, but no treatment of tamoxifen. Examples of ducts and acini are shown.

b. Examples of tdTomato(+) ducts in *TS* mice, as indicated by H&E staining, IHC for tdTomato (counterstain: hematoxylin), and TdTomato(+) immunofluorescence for Amylase (counterstain: DAPI).

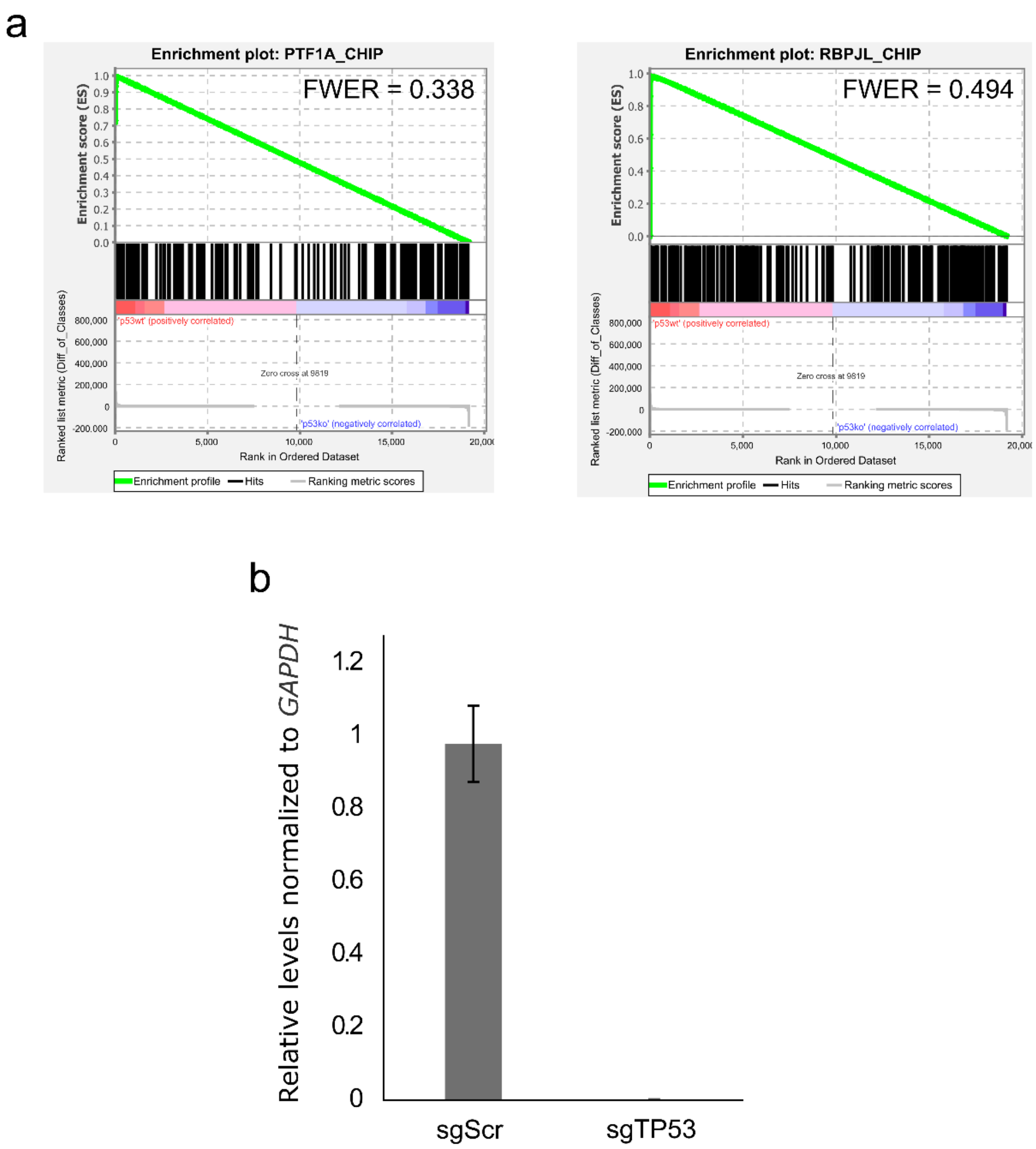


**Figure S4:**

a. Gene set enrichment analysis (GSEA) using RNA-Seq derived from p53 proficient and deficienct pancreatic cells, cross-compared with ChIP-Seq datasets form PTF1A (left) and RBPJL (right).

b. RT-qPCR assessment of CRISPRi knockdown of p53.

**Table S1:**

Sequences of primers used in qPCRs.

| Human; | | |
|---|---|---|
| Gene | Fwd (5' - 3') | Rvs (5' - 3') |
| *TP53* | GCAGTCAGATCCTAGCGTCG | TTTTCAGGAAGTAGTTTCCATAGGT |
| *BHLHA15* | CGGATCCCCAGCTCCAAG | GTTCTTGGTCTTCATGGCCC |
| *CDKN1A* | AGAGGCTGGTGGCTATTTTGT | TCTGACATGGCGCCTGAAAAC |
| *GAPDH* | ATGAGAAGTATGACAACAGCCTCAAGAT | ATGAGTCCTTCCACGATACCAAAGTT |

| Mouse; | | |
|---|---|---|
| Gene | Fwd (5' - 3') | Rvs (5' - 3') |
| *Trp53* | CGGGAAATAGAGACGCTGAG | CCCGAGAAGCCACAGATAAG |
| *Bhlha15* | TCGAATCCCCAGTTGGAAGG | TAGCTCCAGGCTGGTTTTCC |
| *Cdkn1a* | CAGCAGAATAAAAGGTGCCACA | GGAACAGGTCGGACATCACC |
| *Gapdh* | AGGTCGGTGTGAACGGATTTG | TGTAGACCATGTAGTTGAGGTCA |
